## Supplementary material for "PIF transcriptional regulators are required for rhythmic stomatal movements": Rovira et al_SI

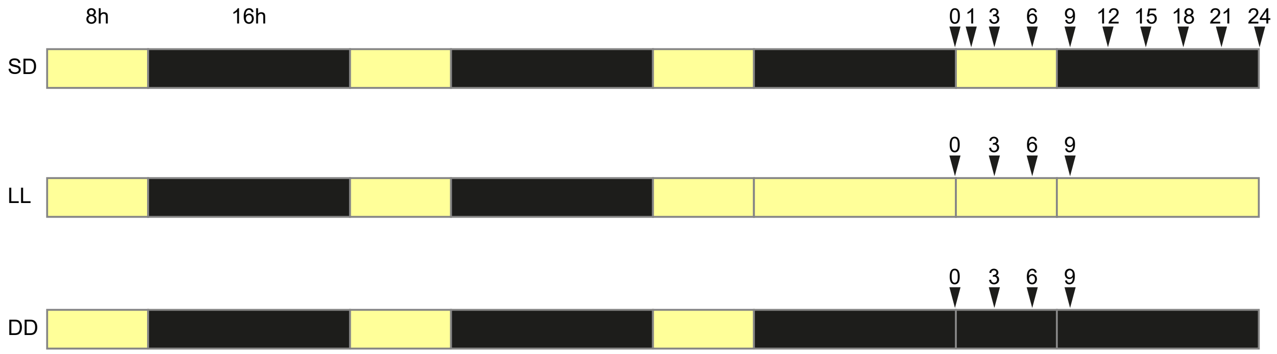


**Fig. S1. Schematic diagram of the SD, LL, and DD growth regimes used.** After vernalization, plates were transferred at the beginning of the day to short days (SD) (8h light + 16 h dark) for seedlings to germinate. Sampling in SD took place in the dark during the end of the third day (ZT=0), and in the light (ZT=1, 3, 6) or in the dark (ZT=9, 12, 15, 18, 21, 24) during the fourth day of growth. Continuous light (LL) samples correspond to time points taken in seedlings entrained in SD and then transferred to continuous light from the third night onward. Continuous dark (DD) samples correspond to time points taken in seedlings entrained in SD and then transferred to continuous dark from the third night onward. Yellow and black rectangles represent light and dark, respectively, and arrow heads indicate sampling time points.

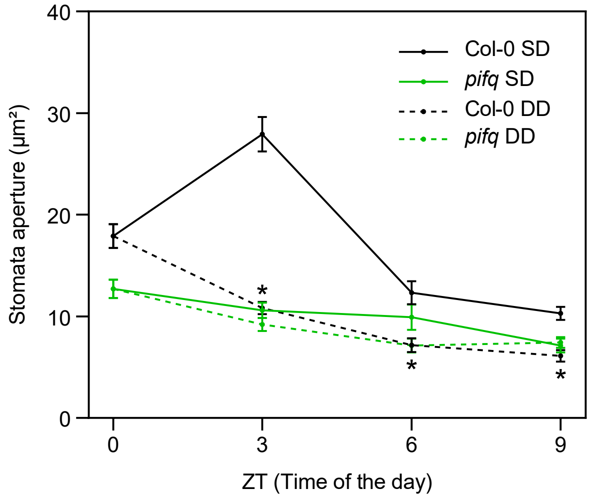


**Fig. S2. Time course analysis of stomata aperture in Col-0 and *pifq* grown under DD.** Seedlings were grown under short day (SD) conditions for 2 days, then at ZT8 of the third day they were either kept under SD or transferred to continuous dark (DD) (see SI Fig. 1 for a diagram of light treatments). Stomata pore measurements were performed during the fourth day at ZT0, 3, 6 and 9h and expressed as area. Time points represent mean values ± SE. n=40. Statistical differences relative to Col-0 for each time point and condition are indicated by an asterisk (Mann-Whitney test. $P<0.05$).

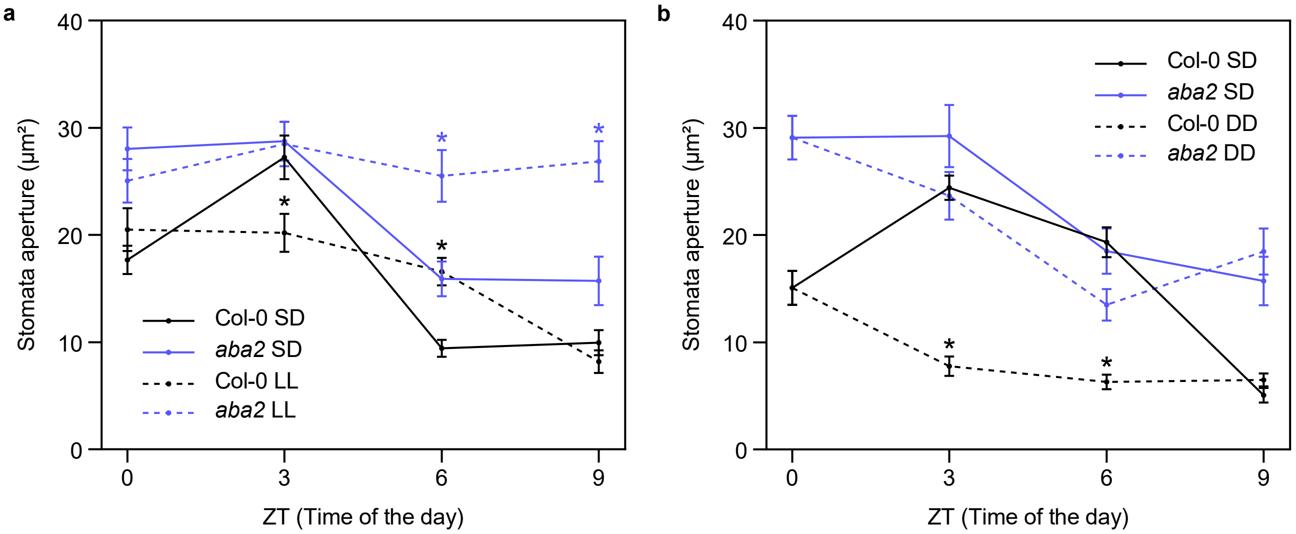


**Fig. S3. Time course analysis of stomata aperture in Col-0 and *aba2* grown under LL or DD.** Seedlings were grown under short days (SD) for 2 days, then at ZT8 of the third day they were either kept under SD or transferred to (a) continuous light (LL) or (b) continuous dark (DD). Stomata pore measurements were performed during the fourth day at ZT0, 3, 6 and 9h and expressed as area. Time points represent mean values ± SE. n=40. (a, b) Statistical differences relative to Col-0 for each time point and condition are indicated by an asterisk (Mann-Whitney test. $P<0.05$).

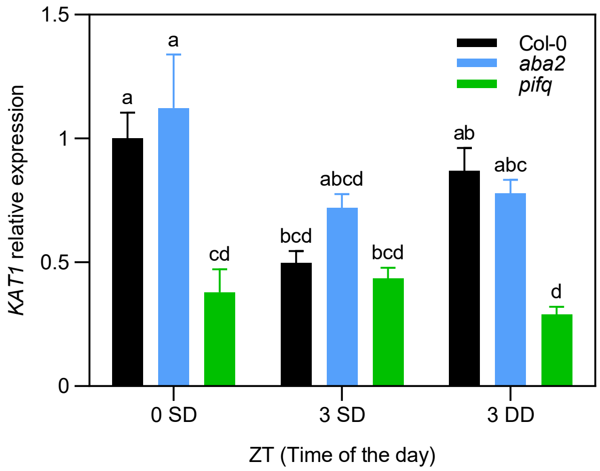


**Fig. S4. *KAT1* expression in Col-0, *pifq*, and *aba2* seedlings under DD compared to SD.** *KAT1* expression in Col-0, *pifq*, and *aba2* seedlings in 3-day-old SD- or DD-grown at ZT0 (common for SD and DD) and ZT3. Data are the means ± SE of biological triplicates (n = 3). Letters denote the statistically significant differences using 2-way Anova followed by posthoc Tukey's test ($P < 0.05$).

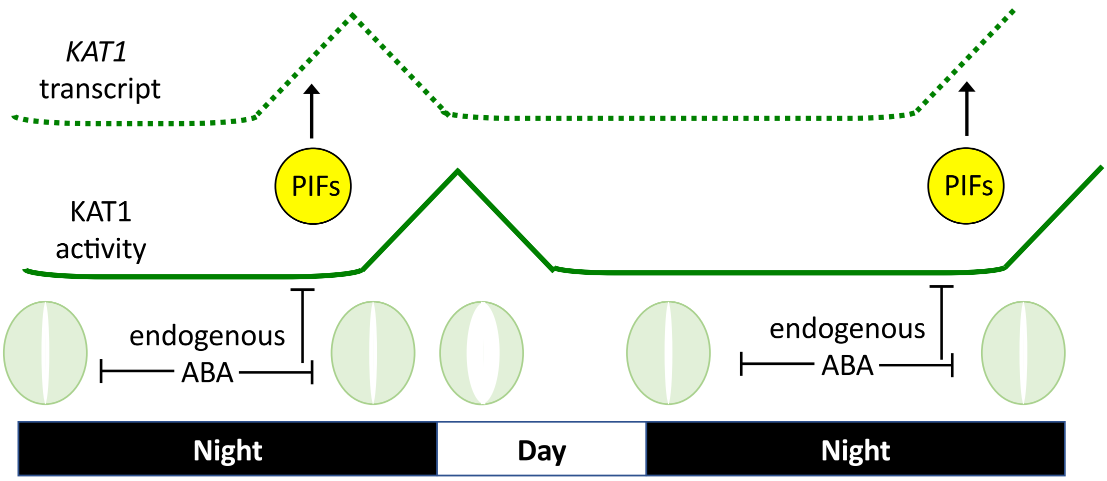


**Fig. S5. Model of PIF and endogenous basal ABA interplay in the diurnal regulation of KAT1 in stomata dynamics.** During the night in short days, PIFs progressively accumulate and induce *KAT1* expression at the end of the night. Endogenous ABA also accumulates at night and prevents activity of KAT1. At dawn, light induces ABA degradation and KAT1 can accumulate and promote stomata opening downstream of light-activated phot1 driving membrane hyperpolarization. Light also triggers phytochrome-mediated PIF degradation, effectively inhibiting *KAT1* overexpression.

**SI Table 2: Primers used for qRT-PCR.**

For each gene analyzed by qRT-PCR the pair of primer sequences is shown. Columns indicate the primer name, the sequence, and the gene amplified. For primers described elsewhere, the reference is indicated.

| Primer | Sequence | Gene |
|---|---|---|
| EMP1123 | GCAATAAGGTACCTTTCGAC | *KAT1* |
| EMP1124 | AAGCCTTGCAAATAGCGAGC | *KAT1* |
| EMP338 | TATCGGATGACGATTCTTCGT | *PP2A*[70] |
| EMP339 | GCTTGGTCGACTATCGGAATG | *PP2A*[70] |

**SI Table 3: Primers used for ChIP-qPCR.**

For each binding region analyzed by ChIP-qPCR the pair of primer sequences is shown. Columns indicate the primer name, the sequence, the gene amplified, and the binding region in the promoter (P1 and P2) or the gene body (P3).

| Primer | Sequence | Gene | Binding region |
|---|---|---|---|
| EMP1152 | GCATGGGAAGTGAAACTCTAAG | *KAT1* | P1 |
| EMP1153 | CGAGTGAGAAGAGAGTTTGGG | *KAT1* | P1 |
| EMP1154 | GCAAGCAATATGTCTTTGTTG | *KAT1* | P2 |
| EMP1155 | CCGACGGGAATGAGAAGTATG | *KAT1* | P2 |
| EMP1178 | CCAAACTTCTCACTTGCAAGTC | *KAT1* | P3 |
| EMP1179 | GATCCATATTGCAGCTCAAGC | *KAT1* | P3 |
